# Reference genome of the ant *Lasius platythorax*

**DOI:** 10.1101/2023.07.18.549505

**Authors:** Barbara Feldmeyer, Nadège Guiglielmoni, Joseph Kirangwa, Florian Menzel, Judit Salces-Ortiz, Rosa Fernández, Elena Buena Atienza, Claudio Ciofi, Maria Angela Diroma, Alessio Iannucci, Chiara Natali, Ann M. Mc Cartney, Olaf Riess, Nicolas Casadei, Ann-Marie Waldvogel

## Abstract

Ants are a highly diversified insect family of the order Hymenoptera, with many fascinating characteristics such as eusociality, chemical communication, farming, or social parasitism. Moreover, ants frequent a wide variety of habitats from dry deserts, grasslands, and savannas to cold temperate forests. The ability of ants to inhabit such diverse habitat ranges demonstrates their adaptability and ecological resilience. However, little is known about the genetic underpinnings of this vast array of traits and their adaptive potential. Here, we generated a high-quality genome assembly for the ant species *Lasius platythorax* using long-read PacBio HiFi in combination with chromatin conformation capture (Hi-C) sequencing. We successfully assembled the genome into 15 chromosome-level scaffolds ranging from 7.9 to 19.2 Mb and encompassing 204.6 Mb out of 235.3 Mb (total assembly), and a BUSCO score of 86% (Hymenoptera_odb10). Comparative genomics between the two sister species will provide insights into the genomic basis of trait differentiation.

## Introduction

Ants inhabit nearly all terrestrial habitats and fulfill multiple ecosystem functions (Elizalde et al. 2020, Wills & Landis 2018), and have many characteristics of interest including social structure, division of labour, morphological and behavioural diversity, and intricate communication systems (Hölldobler and Wilson 1990). The genus *Lasius* (Formicidae: Formicinae) is common and species-rich in many Palearctic habitats and is scientifically significant due to extraordinary behaviours, species interactions, ecological services, but also invasive species dynamics (Hölldobler and Wilson 1990; Cremer et al. 2008; Seifert 2018; Stukalyuk et al. 2020). Some *Lasius* species have evolved a social parasitic lifestyle, where one eusocial species depends on the labour force of another (Buschinger 2009; Savolainen and Vepsalainen 2003). Moreover, fungiculture generates a powerful system for investigating the origin and maintenance of mutualism (Mueller et al. 2005). In ants, fungiculture has evolved at least twice independently: in attines which are members of the Myrmicinae that culture the fungi for food, and in *Lasius* ants that utilise fungi to construct composite nest walls (Maschwitz and Hölldobler 1970; Mueller and Gerardo 2002; Schlick-Steiner et al. 2008). Within the genus *Lasius*, both social parasitism and fungiculture evolved twice (Boudinot, Borowiec, and Prebus 2022; Maruyama et al. 2008). Moreover, *Lasius* ants are highly diverse in their signal and defense chemistry (Akino and Yamaoka 2005; Kern et al. 1997; Wu and Mori 1991; Yamaoka and Akino 2000). Ants of the genus *Lasius* are widely distributed throughout most of the northern hemisphere, from the Indomalaya, Nearctic, Neotropical, Palearctic (Quque and Bles 2021).

*Lasius platythorax* is a small brownish ant, which is closely related to *Lasius niger* and was recognised as a separate species in only 1991 (Seifert 1991). While *L. niger* is among the most common species throughout Central Europe, inhabiting grasslands and park areas, *L. platythorax* inhabits deciduous forests, bogs, and fens. *L. platythorax* prefers habitats that are more humid and less sunny in comparison to its sister species. These sibling ant species can be separated using morphometric characters and ecological parameters (Seifert 2007). In addition to inhabiting different niches, *L. platythorax* also differs behaviourally, displaying a higher level of aggression than *L. niger* (Seifert 2007). Interestingly, the cuticular hydrocarbons of *L. platythorax* and *L. niger* also differ significantly. Cuticular hydrocarbons play an important role in ant communication, particularly in recognition of nestmates, and in waterproofing and defense against pathogens (Lenoir et al. 2009). *L. platythorax* cuticular hydrocarbons are characterised by a high abundance of dimethyl and trimethyl alkanes with relatively high chain lengths (mostly between C33 and C37), and so are longer than those of *L. niger* (Wittke, Baumgart, and Menzel 2022; Baumgart et al. 2022).

To date, genomic research on *Lasius* ants has been limited, with only a single *Lasius* genome available for the black garden ant *L. niger* (Konorov et al. 2017). A broader understanding of genomic diversity, intricate communication systems, and adaptive potential within the *Lasius* genus requires the availability of multiple genome assemblies representing different species. Here, we present a reference genome of *L. platythorax*, generated via a combination of long-read Pacific Biosciences (PacBio) HiFi sequencing and chromatin conformation capture (Hi-C). This genome will provide invaluable opportunities to investigate the genomic basis of ecological, behavioural, and chemical differentiation among closely related *Lasius* species as well as gain insights into the mechanisms driving their adaptive potential and species diversification.

## Materials and Methods

### Sample collection and DNA extraction

*Lasius platythorax* is a small, brownish ant. It is distributed across Central Europe and is typically found in different kinds of woodland, bogs, and fens and avoids urban habitats. The species is monogynous and colonies can contain several thousands of workers (Seifert 1991). One *L. platythorax* colony was sampled in the Lennebergwald, close to Mainz in May 2021. Brood and workers were collected from this single colony for the long-read and Hi-C sequencing effort. DNA was isolated using the Qiagen MagAttract HMW DNA Kit from a single larva.

### RNA extraction

RNA was extracted with the Zymo Direct-zol RNA miniprep kit, from a pool of ten pooled larvae and five pooled workers, according to the manufacturer’s instructions. RNA quality and quantity were determined using the Nanodrop 2000 and stored at -80°C.

### Long-read sequencing

HiFi libraries were prepared with the Express 2.0 Template kit (Pacific Biosciences, Menlo Park, CA, USA) from a single larva and sequenced on a Sequel II/Sequel IIe instrument with 30h movie time.

HiFi reads were generated using SMRT Link (v10; (Pacific Biosciences, Menlo Park, CA, USA) with default parameters.

### Hi-C library and sequencing

Hi-C libraries were prepared from a single larva using Arima Genomics HiC kit, adhering to the manufacturer’s guidelines with specific adjustments, including extended lysis because of the thick cuticle and overnight chromatin digestion. The sample was flash-frozen and stored at -70ºC before processing. DNA fragmentation was performed using a Bioruptor Pico (Diagenode), and libraries were generated with the KAPA Hyper Prep kit (P/N: KK8504). Library concentration and fragment size distribution were assessed using a Qubit Fluorometer (Thermo Fisher Scientific) and a Bioanalyzer with the DNA 1000 Kit (Agilent Technologies), respectively. Arima Hi-C libraries were pooled, denatured, and loaded for paired-end sequencing on Illumina NovaSeq 6000 system (NovaSeq 6000 S1 Reagent Kit v1.5, 300 cycles).

### RNA sequencing

Library preparation for full-length mRNASeq was performed using the NEB Ultra II Directional RNA Library Prep with NEBNext Poly(A) mRNA Magenetic Isolation Module and 100ng total RNA as starting material. Sequencing was performed on an Illumina NovaSeq 6000 device with 2x150bp paired-end sequencing protocol and >50M reads per sample.

### Genome assembly

PacBio HiFi reads were assembled using wtdbg2 v2.5 (Ruan and Li 2020) with parameters - x ccs -g 200m. Hi-C reads were trimmed using TrimGalore v0.6.10 (Krueger et al. 2023), mapped to the draft assembly using bowtie2 v2.2.5 (Langmead and Salzberg 2012) and preprocessed using hicstuff v3.1.5 (Matthey-Doret et al. 2020) with parameters -e DpnII,HinfI -m iterative. The assembly was then scaffolded based on Hi-C contacts using instaGRAAL v0.1.6 (Baudry et al. 2020) no-opengl branch with parameters --level 4 --cycles 50 and curated using the module instagraal-polish with parameters -m polishing -j NNNNNNNNNN. Gaps were filled using TGS-GapCloser v1.1.1 (Xu et al. 2020) with parameters --ne --minmap_arg ‘-x map-hifi’. PacBio HiFi reads were mapped using minimap2 v2.24 (Li 2018) with parameters --secondary=no --MD -ax map-hifi and the output was provided to HyPo v1.0.3 (Kundu, Casey, and Sung 2019) for polishing with parameters -k ccs -s 200m -c 70. *k*-mer completeness was assessed with KAT comp v2.4.2 (Mapleson et al. 2017) and orthologue completeness was evaluated using BUSCO v5.0.0 (Manni et al. 2021) against the Hymenoptera odb10 and the Insecta odb10 lineages in genome mode. Meryl v1.3 and Merqury v1.3 (Rhie et al. 2020) were run with default parameters using the PacBio HiFi reads. The scaffolds were aligned against the nucleotide database using the Basic Local Alignment Search Tool (BLAST) v2.13.0 (Altschul et al. 1990) with parameters -outfmt “6 qseqid staxids bitscore std sscinames scomnames” -max_hsps 1 -evalue 1e-25. The PacBio HiFi reads were mapped against the scaffolds using minimap2 v2.24 with parameter -ax map-hifi and the alignment was sorted using SAMtools v1.17. The outputs of minimap2, BLAST, and BUSCO were provided as input to BlobTools2 v4.1.5. The Hi-C contact map of the main scaffolds was built using hicstuff as previously described and visualized using the module hicstuff view with parameter -b 500.

### Repeat masking and gene annotation

We first used RepeatModeler v2.0.4 (Flynn et al. 2020) to generate a custom repeat library. We then ran two rounds of RepeatMasker v4.1.4 (Tarailo-Graovac and Chen 2009), the first round with the custom repeat library, search engine “ncbi”. The second round of RepeatMasker was run with -species “arthropoda”. The RepeatMasker output tables were merged to summarize the abundance or RE categories with the ProcessRepeats function of RepeatMasker. We used HiSat2 (Kim et al. 2019) to map transcriptome reads to the genome and used the .bam file as RNA-seq evidence. BRAKER3 (Gabriel et al. 2023) was used to annotate the softmasked genome using RNA-seq evidence, OrhtoDB11 protein evidence (Kuznetsov et al. 2023) as well as setting the -AUGUSTUS_ab_initio option. BUSCO v5.2.2 (Manni et al. 2021) against the Hymenoptera odb10 and the Insecta odb10 lineages in protein mode was used to assess the completeness of the annotation.

## Results

The genome of *Lasius platythorax* was assembled from 25.0 Gb of PacBio HiFi reads with an N50 of 11.3 kb and 273 million pairs of Hi-C reads. Initial assembly with wtdbg2 yielded 3,986 contigs with a total size of 235.3 Mb and an N50 of 106 kb (Table 1). 79% of the Hi-C reads were mapped to this draft assembly. Hi-C scaffolding using instaGRAAL resulted in 1,374 scaffolds with 15 chromosome-level scaffolds ranging from 7.9 to 19.2 Mb and encompassing 204.6 Mb out of 235.3 Mb (total assembly, Figure 1). Gap filling and polishing closed 204 gaps to reach 3,332 gaps and increased the assembly size to 235.8 Mb. The final 15 chromosome-level scaffolds have BUSCO scores of 85.6% (0.4% duplicated features) against Hymenoptera odb10 and 90.1% (0.8% duplicated features) against Insecta odb10. *k*-mer analysis does not show uncollapsed haplotypes (Figure 2) and the QV score of 43.7 indicates high overall accuracy. No contaminants were identified (Figure 3).

**Table 1:** Genome features of *Lasius platythorax* including the mitochondrial and plastid genomes.

|  |  |
| --- | --- |
| Assembly ID | *L. platythorax* genome v1 |
| Total length (bp) | 235,769,877 |
| No. of contigs | 3,986 |
| No. of scaffolds | 1,374 |
| Longest scaffold (bp) | 19,197,288 |
| N50 (kb) | 15,532,059 |
| L50 | 7 |
| GC content (%) | 36.12 |

**Figure 1:**
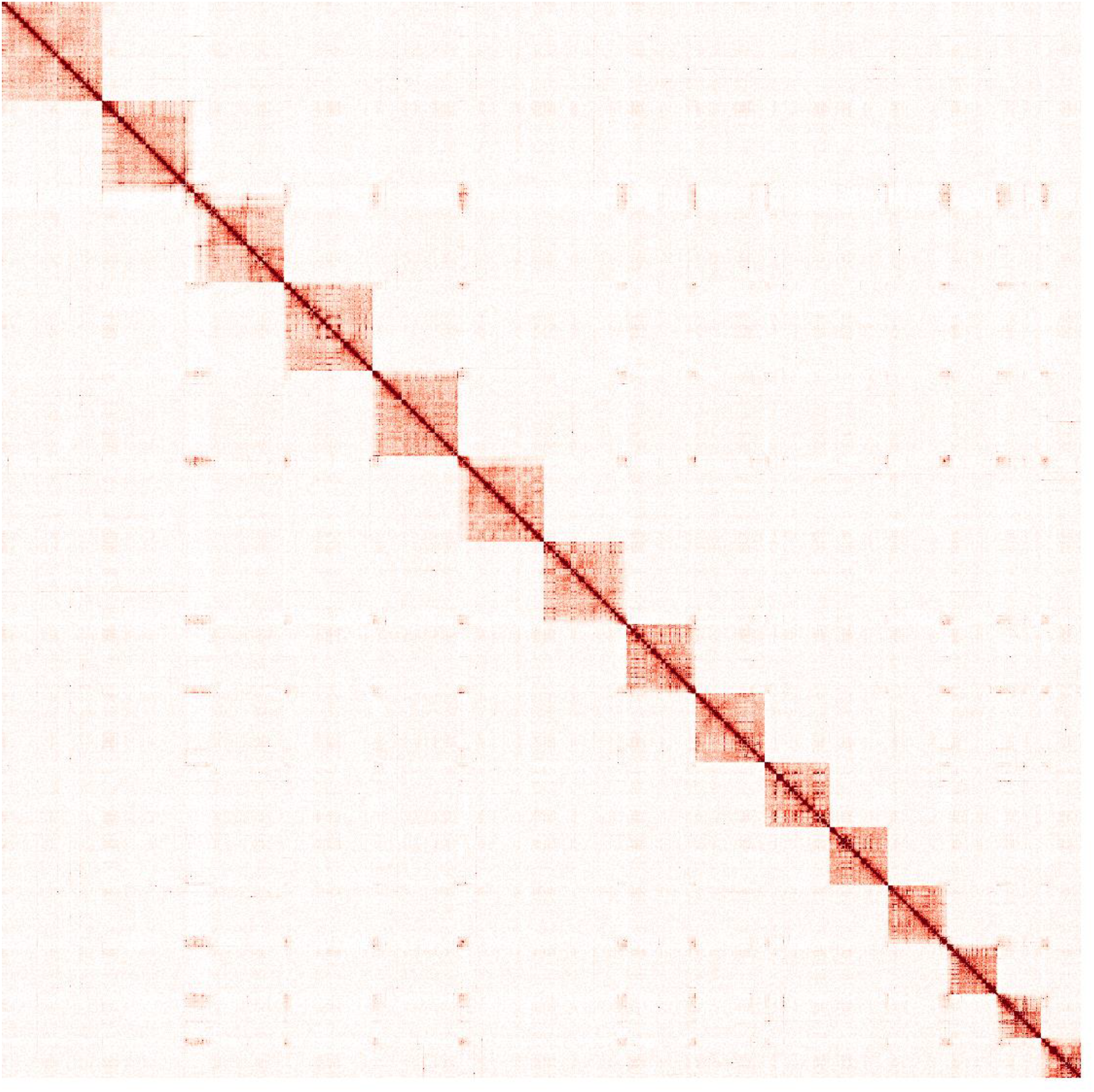
Hi-C contact map of 15 chromosome-level scaffolds.

**Figure 2.**
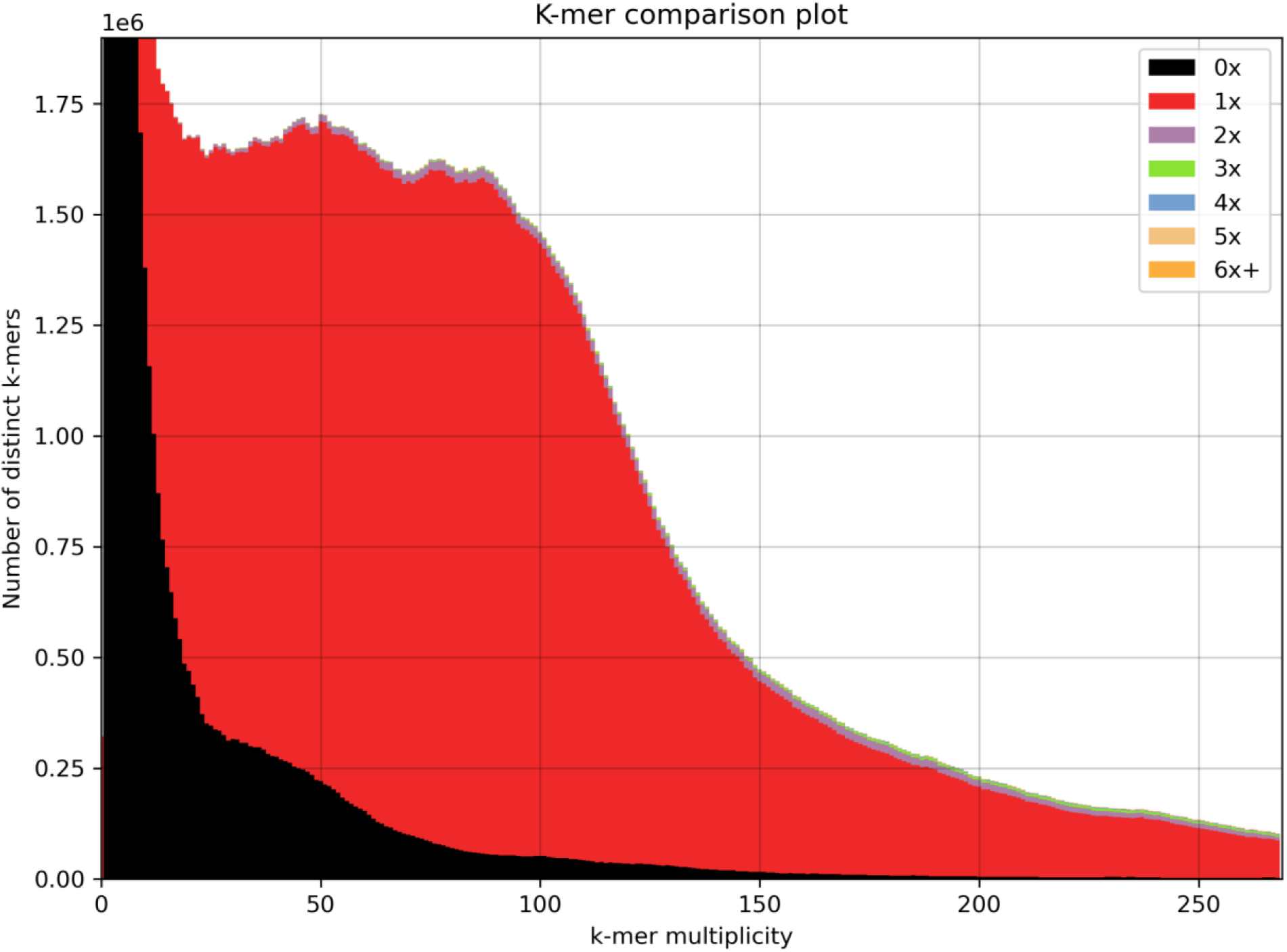
*k*-mer comparison of the HiFi reads v. the final scaffolds.

**Figure 3.**
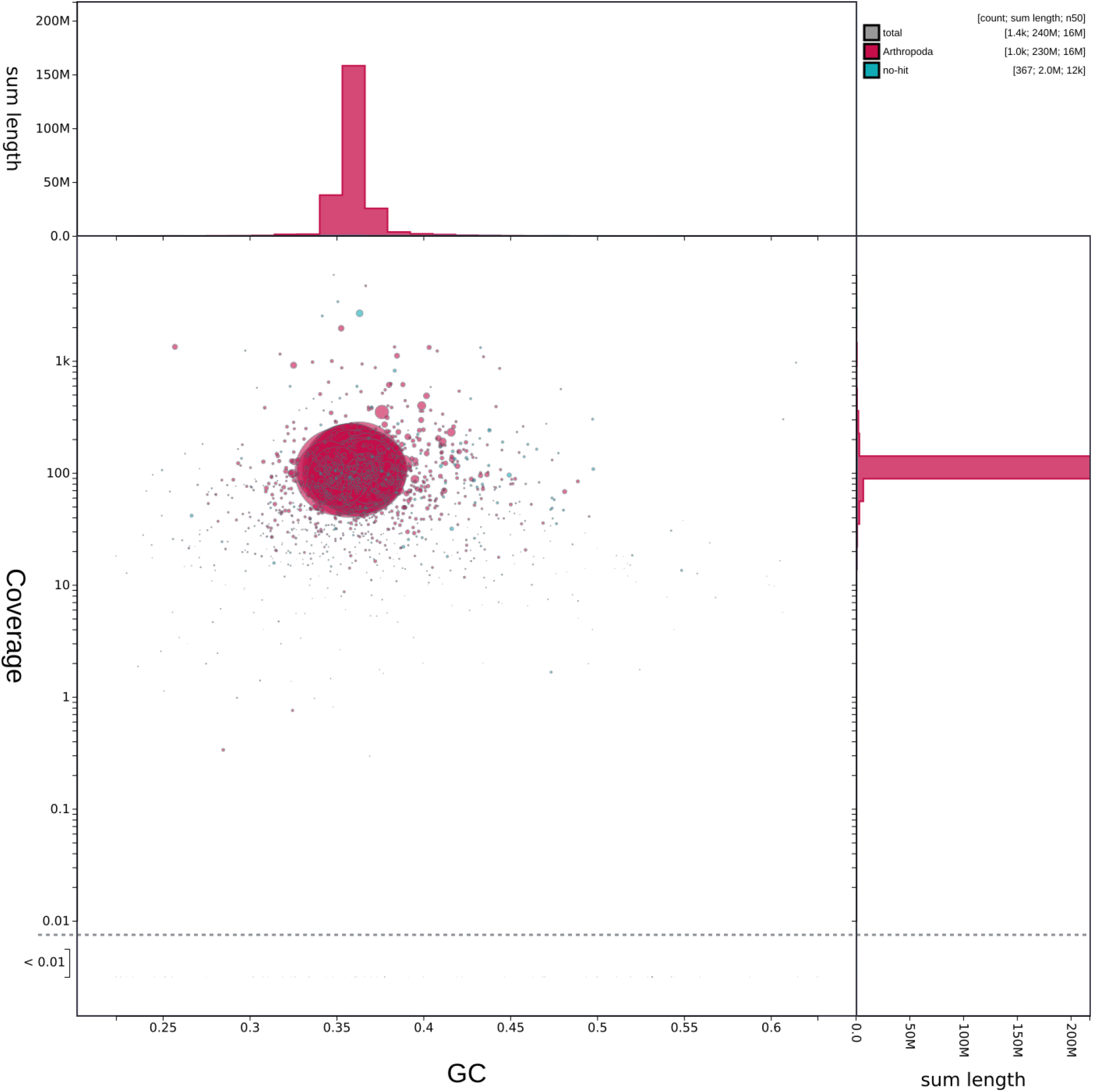
BlobTools analysis of the final scaffolds. The sequences were either flagged as “Arthropoda” or “no hit”, and do not exhibit drastic divergence in coverage or GC content.

Approximately 23% of the genome assembly consists of repetitive elements, which is comparable to other ant genomes (Hartke et al. 2019). These 23% were mainly annotated as simple repeats (3.23%) and interspersed repeats (18.72%) (Table 2). Within the latter, the amount of retroelements (6.02%), DNA transposons (5.88%), and unclassified repeats (6.83%) is comparable. The annotation with BRAKER3 (Gabriel et al. 2023) using a combination of RNA-seq and protein evidence as well as ab-initio gene prediction resulted in the annotation of 17,976 genes. A total of 1208 (88%) complete gene models among 1367 conserved orthologs were identified by BUSCO (Table 2).

**Table 2:** Annotation features of the *L. platythorax* genome.

| Genome annotation |  |
| --- | --- |
| Repeat content (%) | 23.29 |
| Retrotransposons | 6.02 |
| DNA transposons | 5.88 |
| Simple repeats | 3.23 |
| Unclassified | 6.83 |
| No. gene models | 17,976 |
| BUSCO (insecta_odb10) | C:88.4%[S:71.8%,D:16.6%],F:1.0%,M:10.6%<br>n:1367 |

## Conclusion

This newly generated *L. platythorax* genome will provide valuable opportunities to investigate the genomic basis of niche, behavioural and chemical differentiation among closely related *Lasius* species and gain insights into the mechanisms driving their adaptive potential and species diversification.

## Data availability

The data are part of the ERGA pilot project. Long-read PacBio HiFi data are accessible via BioProject: PRJNA1054215, and BioSample: SRX23153489. Illumina RNAseq and Illumina HiC raw sequences, as well as the assembly and annotation were uploaded to ENA with the accession: PRJEB72067. TolIDs=iyLasPlat10-17:001-008.

## Acknowledgments

This genome project is part of the DeRGA pilot study and we acknowledge coordination and support by Dr. Astrid Böhne and Dr. Philipp Schiffer. We thank Stephan Ossowski for bioinformatic support with the HiFi data. We acknowledge collaboration with the ERGA Pilot Project Coordinators, particularly Dr. Alice Mouton and Dr. Giulio Formenti. We furthermore acknowledge and thank all supplier partners that have kindly donated kits, reagents to the ERGA pilot Library Preparation Hubs to support species to produce the generation of high-quality genomes and annotations. This support has been key to embedding a culture of diversity, equity, inclusion, and justice in the Pilot Project. Specifically we want to thank Arima Genomics, Integrated DNA Technologies (IDT), Agilent Technologies, Fisher Scientific Spain and Illumina Inc.

